# Human gut flagellome profiling using FlaPro reveals TLR5-related phenotype-specific alterations in IBD

**DOI:** 10.1101/2025.06.30.660334

**Authors:** Anna A. Bogdanova, Andrea Borbón-Garciá, Ruth E. Ley, Alexander V. Tyakht

## Abstract

**Background:** Flagellin is the protein monomer of the bacterial flagellum, which confers motility, allowing bacteria to reach their favored niches. Flagellin is highly conserved across bacterial species and thus the target of the innate immune receptor Toll-like receptor 5 (TLR5). In the gut, bacterial flagellin agonizes human TLR5, triggering a pro-inflammatory response. However, the ability to bind and activate TLR5 varies considerably between different flagellins, suggesting that the composition of an individual’s flagellin repertoire – the flagellome – may mediate the inflammatory response to the microbiome, with relevance to inflammatory bowel diseases. However, to date, methods to assess the inflammatory potential of a flagellome are lacking.

**Methods:** We constructed a curated database of human gut microbiome-derived flagellins. To predict the inflammatory potential of the flagellome by sorting flagellins into either “stimulatory” (strong TLR5 agonists) or “silent” (weak TLR5 agonists), we trained a machine learning model on experimentally characterized flagellins with known binding and stimulatory activities. The FlaPro pipeline was implemented using the Snakemake workflow engine for high-throughput analysis and is available at https://github.com/leylabmpi/FlaPro. A publicly available multi-omics dataset from an inflammatory bowel disease (IBD) cohort was used to explore associations between flagellome features and clinical status.

**Findings:** FlaPro enables robust profiling of the human gut flagellome from metagenomic and metatranscriptomic data. Analysis of the IBD datasets revealed a depletion of flagellome diversity and a reduced silent-to-stimulatory flagellin abundance ratio in Crohn’s disease and ulcerative colitis, observed at both the genomic and transcriptional levels. Multiple condition-specific alterations were identified at the level of individual flagellin clusters.

**Interpretation:** These findings indicate that IBD is associated with distinct alterations in the gut flagellome, particularly in relation to TLR5 recognition. Flagellome features represent a functionally interpretable class of microbiome-derived markers with potential utility in microbiome-wide association studies in the context of human health and disease.

**Funding:** This work was supported by the Max Planck Society and the European Research Council (ERC) under the European Union’s Horizon 2020 research and innovation programme Grant agreement ID: 101142834 (ERC Advanced Grant SilentFlame).

## Introduction

The gut microbiota plays an essential role in human physiology, contributing to digestion, interacting with the immune system and influencing distal organ systems. As a symbiotic community, the human gut microbiota is generally more commensal or beneficial than harmful; however, an imbalance in microbial composition can compromise host health due to the pathogenic potential of specific taxa. Maintaining the balance by eliminating harmful bacteria while preserving commensal and beneficial ones is therefore crucial. One mechanism by which the immune system recognizes bacteria is through pattern recognizing receptors (PRRs) such as Toll-like receptor 5 (TLR5), which detects flagellins and initiates immune response.

Flagellin is the protein structural constituent of prokaryotic flagella, which act as locomotive nanomachines enabling bacterial movement. Due to its role in motility and surface exposure, flagellin serves as a key target for immune recognition. The structure of flagellins comprises two conserved domains, D0 and D1, at both the C-terminal and N-terminal ends, connected by a hypervariable region (HVR) containing domains D2 and D3. Initial models based on the FliC flagellin of the pathogen *Salmonella* Typhimurium proposed the interaction between flagellin and human TLR5 involves three main steps: (i) TLR5 detects a conserved region in flagellins, triggering proinflammatory signaling; (ii) this leads to the recruitment of MyD88 and activates downstream cascades culminating in the activation of NF-κB; (iii) NF-κB translocates to the nucleus and promotes transcription of pro-inflammatory cytokines (Yiu, Dorweiler, and Woo 2017). Subsequent studies have shown that *Helicobacter pylori* flagellins fail to bind and therefore evade TLR5-mediated detection (Gewirtz et al. 2004). This prevailing model that flagellin-TLR5 binding led to activation failed to explain the poor agonism of flagellins produced by common gut commensal bacteria, such as the Lachnospiraceae.

Flagellins with poor agonism elicit a TLR5 response at orders of magnitude higher concentrations than the canonical stimulatory flagellin FliC. Clasen et al. observed a so-called “silent” behavior, whereby flagellins bind TLR5 strongly at the D1 domain similarly to stimulatory flagellins, but fail to elicit a robust response. This poor agonism is due to the lack of a previously undescribed allosteric binding site on the D0 domain necessary for signalling (Clasen et al. 2023). Based on these observations, flagellins tested for their TLR5 binding and activation could be classified into “stimulatory”, “evasive” and “silent” (Clasen et al. 2023). However, the features that confer these phenotypes remain to be fully explored for the purpose of classifying novel flagellins.

High-throughput metagenomics enables deep characterization of a human microbiome sample by capturing not just the taxonomic composition but also the functional potential. Metatranscriptomics extends this by revealing the actual expression of encoded functions. Associative studies of large cohorts have helped identify both shared and condition-specific microbial signatures of various human diseases. Given the immense dimensionality of such feature space – achieving the orders of hundreds of millions of features in gene catalogs (Almeida et al. 2021) – practical analysis typically requires reducing it via feature aggregation or targeting specific functionally relevant gene classes (e.g. carbohydrate-active enzymes, antibiotic resistance genes and virulence factors, or pathway-based metabolic phenotypes) (Iablokov et al. 2020). Although flagellins comprise a relatively compact family within the gut microbiome’s gene repertoire, considering the variability of their TLR5 interaction phenotypes it is promising to: i) characterize the complete pool of flagellins within individual samples (hereafter, the flagellome); and ii) explore associations of flagellome with host health, disease states or other factors.

Inflammatory bowel diseases (IBD) – ulcerative colitis (UC) and Crohn’s disease (CD) – are marked by abnormal immune responses to commensal gut microbiome. Bacterial flagellins have been implicated in the onset and progression of these pathologies. There is a broad consensus that immune responses to flagellins play an important role in the CD pathogenesis. In particular, elevated levels of anti-flagellin antibodies (IgA and IgG) are consistently observed in CD patients and employed as diagnostic markers (Lodes et al. 2004; Alexander et al. 2021; Shen et al. 2008). These responses are typically mediated by flagellin-specific CD4^+^ T cells activated via TLR5 signalling pathways, driving the chronic intestinal inflammation. Of note, a Lachnospiraceae flagellin CBir1, the immunodominant antigen recognized by colitic mice and by approximately half of patients with CD (Targan et al. 2005), has been categorized as a “silent” flagellin (Clasen et al. 2023). While the role of flagellin-specific immune responses in UC is less well defined, the presence of anti-flagellin antibodies in UC patients, although at lower prevalence (Cook et al. 2020), suggests a possible involvement in disease pathogenesis as well. Together, these observations suggest that silent flagellins may interact differently from stimulatory flagellins with adaptive immunity in IBD. However, to date, methods to sort flagellins from flagellomes derived from metagenomes and metatranscriptomes are lacking.

We developed FlaPro – a modular workflow for high-throughput quantification of the flagellome in human gut metagenomes and metatranscriptomes. Using experimentally derived data, we constructed an TLR5 interaction phenotypes predictive model and a reference database of flagellins annotated with these predictions. We then performed a flagellome-wide association study of IBD gut microbiome based on a published dataset (Lloyd-Price et al. 2019).

## Materials and methods

### Flagellin reference database

We obtained the initial human gut flagellin gene dataset (5,131 sequences) during our previous study (Clasen et al. 2023) by mapping a collection comprising over 33,000 flagellin protein sequences (Hu and Reeves 2020) against human gut metagenomic data, with taxonomic classification for each flagellin obtained using the taxonomizR 0.10.6 R package (Sherrill-Mix, n.d.). We curated the initial set by filtering out the sequences assigned to the taxa not previously reported in the human gut microbiota (Dai et al. 2022), with the exception of opportunists or pathogens and transient species (i.e. food-associated species). This step resulted in the final dataset of 2,543 flagellin gene sequences.

### FlaPro model for predicting TLR5-related phenotypes

To functionally annotate the flagellins as either “stimulatory” or “silent” with respect to TLR5, we leveraged experimentally derived TLR5 binding and activation data available for 122 flagellins (Clasen et al. 2023), of which 92 had been classified as either “stimulatory” and “silent”. These included the silent flagellin RhFlaB of *Roseburia hominis*, and the stimulatory StFliC from *Salmonella* Typhimurium (Clasen et al. 2023). As this annotated set represented only 3.5% of the reference database, we trained a random forest model to predict TLR5-related phenotypes for the remaining sequences. Model training employed 10-fold nested cross-validation (70% training, 30% testing, random sampling) using the nestedcv 0.7.9 R package (Lewis et al. 2023).

The model incorporated 51 features, including: (i) alignment metrics to the cD0 domains of RhFlaB and StFliC (n = 6); (ii) presence of a basic residue (R/K) at position RhFlaB 478 previously linked to stimulation potential (n = 1); (iii) charge and hydrophobicity scores of 21 residues involved in TLR5 interaction (Song et al. 2017; Yoon et al. 2012; Smith et al. 2003; Ivičak-Kocjan et al. 2018; Forstnerič et al. 2016) (n = 42); and (iv) structural similarity to RhFlaB assessed with Foldseek (van Kempen et al. 2024) using RMSD and TM-score metrics (n = 2) (Fig. 1 A).

**Figure 1.**
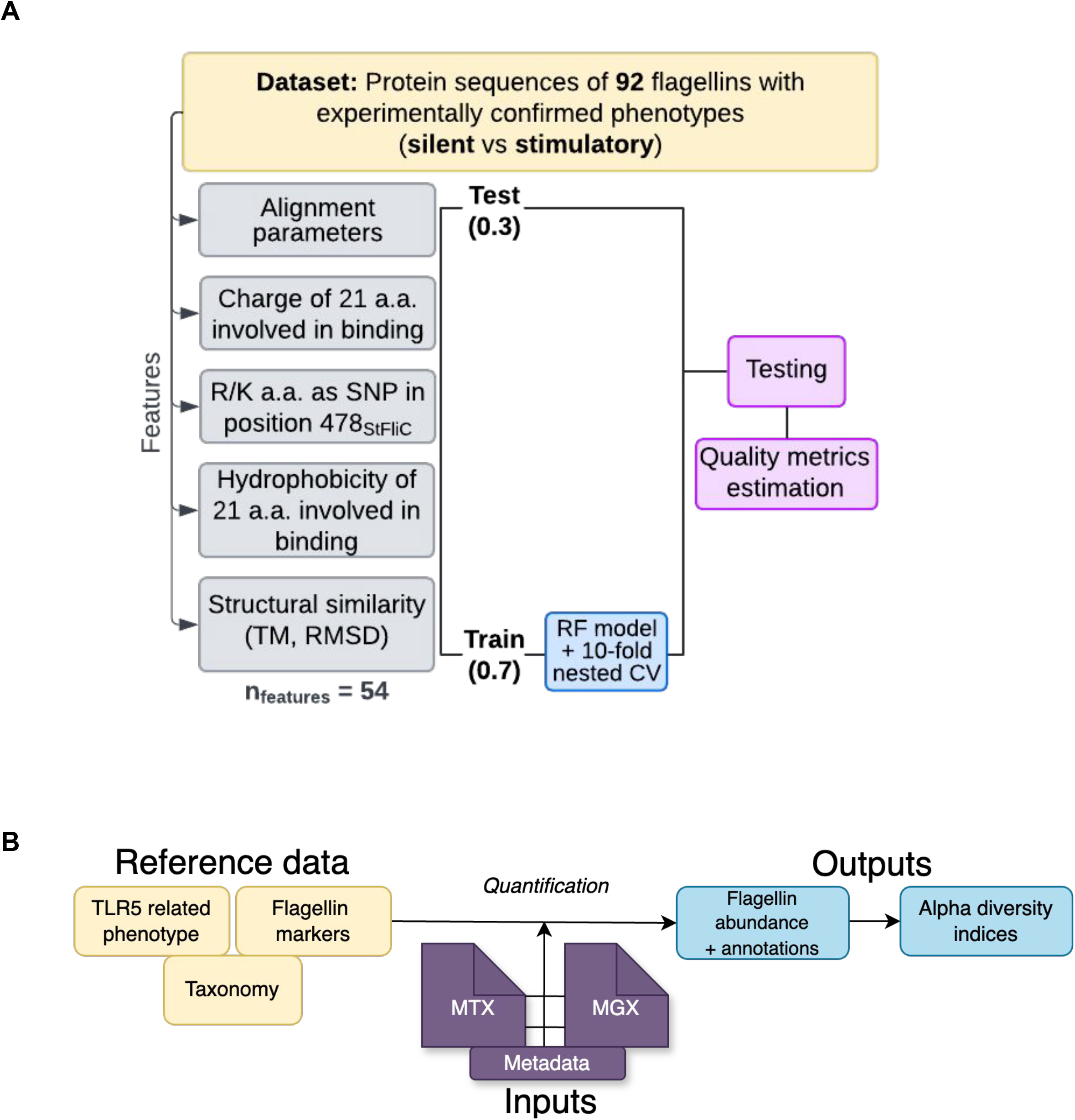
Overview of the FlaPro classification model and workflow. A) Scheme of the classification model showing key components: sample size, phenotype labels, extracted features, and the machine learning algorithm. B) Conceptual diagram of the FlaPro workflow. Using the curated reference data, the user-provided input (metagenomic or metatranscriptomic datasets, MGX or MTX) are processed through a modular pipeline.

Since evasive flagellins were underrepresented in the experimental data, we did not include them into the training set. Thus, the model does not explicitly predict the evasive phenotype. Instead, such sequences are classified into either the stimulatory or (most often) silent categories. For the purpose of this analysis, evasive flagellins are treated as a subclass within the silent category.

Classification confidence score represents the signed distance from the decision boundary (P = 0.5) in probability space. Negative values indicate confidence toward stimulatory phenotype classification, positive values indicate confidence toward silent phenotype classification. Values near zero represent ambiguous cases where classification probability approaches the decision threshold. We used the score to assess the model decision confidence towards false predictions. The scoring formula was defined as:

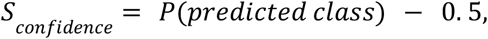

where the sign of the score indicates the direction of the predicted phenotype.

### Workflow for flagellome profiling

The FlaPro pipeline processes gut metagenomes to quantify the reads belonging to flagellins annotated as stimulatory or silent. We implemented the analysis using the Snakemake workflow management system (Mölder et al. 2021), organizing the pipeline into modular components encompassing reference input handling and flagellin quantification. Input data consist of FASTQ or FASTA files derived from metagenomic or metatranscriptomic sequencing. In addition to user-provided microbiome datasets, the workflow requires three reference inputs: (i) the human gut flagellin marker sequence database, (ii) taxonomical annotations, and (iii) functional annotations of the flagellins. The pipeline processes these inputs through a set of modular steps, which can be selectively activated depending on the analysis objectives. Key steps include quantification of flagellins and assignment of taxonomical and functional labels (Fig. 1 B).

For quantifying the presence of specific flagellin sequences, we employed the ShortBRED tool, which consists of two main modules: ShortBRED Identify and ShortBRED Quantify (Kaminski et al. 2015). The Identify module clusters protein sequences based on homology and selects unique marker sequences for each cluster. These markers are subsequently used by Quantify to detect and quantify corresponding sequences in meta’omic datasets. From the curated set of 2,543 flagellin sequences, ShortBRED generated 1,322 clusters. Although cluster-level analysis reduces resolution, it confers a key advantage: flagellins with sequence variation that would otherwise evade exact database matching can still be detected within clusters, allowing broader coverage and improved sensitivity compared to traditional methods. We implemented a modification of the ShortBRED software by replacing USEARCH with DIAMOND2 (Buchfink, Reuter, and Drost 2021) aligner, which is known for higher performance in large-scale analyses.

The quantification module then uses the marker database to analyze either metagenomic (MGX) or metatranscriptomic (MTX) sequencing data. The output includes a table with columns for Flagellin ID, Counts, Hits, and TotalLengthMarker. When <50% of a cluster’s marker sequences are matched by reads from a given sample, the abundance is deemed unreliable by ShortBRED and set to zero; otherwise, abundance is normalized to RPKM (reads per kilobase of reference sequence per million sample reads). To retain the discrete character of the data for downstream statistical analyses, we store original read counts – filtered by the 50% detection threshold – as “real counts”. In combination with taxonomic and predicted phenotype annotations, the resulting relative abundance matrix across multiple samples can be used to assess diversity (with alpha diversity integrated into the pipeline) and perform differential abundance analyses. The pipeline is available at: https://github.com/leylabmpi/FlaPro.

### IBD gut microbiome data

To explore the alterations of flagellome with disease, particularly in IBD, we applied FlaPro to a publicly available stool microbiome dataset from IBD patients and healthy subjects (n = 103) that included both metagenomic (MGX) and metatranscriptomic (MTX) data across multiple time points (Lloyd-Price et al. 2019). The cohort consisted of 26 healthy controls, 28 individuals with ulcerative colitis (UC), and 50 with Crohn’s disease (CD) (Suppl. Fig. 1 A).

### Associations between flagellome features and clinical status

Secondary analysis of the flagellome profiles was implemented as an R-based Jupyter notebook. To enhance reproducibility and facilitate code maintenance, we developed a modular system that assembles project notebooks from reusable, version-controlled code blocks, with reverse synchronization via Git. The analysis included two main approaches: component-based and compositional-aware, each offering distinct perspectives on flagellome variation.

In the component-based analysis, we focused on inter-individual variability in the proportion of flagellated bacteria – and thus flagellin abundance – in a sample. In accord, for both metagenomic (MGX) and metatranscriptomic (MTX) data, relative abundance of each flagellin cluster in a sample was calculated by dividing the cluster’s read counts by the total number of reads in the sample, scaled by a factor of 10^8^. Major flagellin clusters were defined as those with ≥30% prevalence across the samples. Leveraging paired MGX and MTX datasets of the IBD dataset (Lloyd-Price et al. 2019), we quantified the transcriptional activity of each flagellin cluster by computing the MTX/MGX ratio – defined as the MTX-derived relative abundance divided by its MGX-derived counterpart (limited to the clusters with non-zero MGX abundance). For each of the three feature sets (MGX, MTX, and MTX/MGX ratio), we calculated per-sample flagellome alpha diversity indices (Chao1, Shannon, observed features) and the total flagellome relative abundance. These metrics were also stratified by flagellin class (stimulatory, silent, mixed, not defined); and the silent-to-stimulatory abundance ratio was computed. Pairwise dissimilarity of flagellome profiles was assessed using Euclidean distance. Differential abundance analysis for the IBD dataset was conducted on major flagellin clusters following log(x + 1) normalization, using a linear mixed-effects model (lmer function from the lme4 R package), with clinical group and age as fixed effects and subject ID as a random effect. Comparisons included both two-group tests (healthy controls vs. UC; healthy controls vs. CD) and a three-group test using a composite “disease score” variable (0 = HC, 1 = UC, 2 = CD) to reflect increasing disease severity. Multiple testing correction was applied using the Benjamini–Hochberg procedure, with a significance threshold of FDR < 0.05.

In the compositional-aware analysis (applied to both MGX and MTX data), flagellome profiles were treated as compositional data and analyzed using the Nearest Balance (NB). NB identifies associations with a factor (such as disease) by evaluating ratios of the features rather than their individual abundance values and aggregating them into a single fraction (balance) (Odintsova, Klimenko, and Tyakht 2022).

As required by the NB method, we first excluded samples with insufficient total flagellin abundance (<100 reads), then selected the most prevalent flagellin clusters using the same prevalence threshold as in the component-based approach, followed by a second filtering step retaining only clusters with total counts >30. General association between the factor (clinical status) and flagellome composition was tested using PERMANOVA (adonis2 function from vegan R package; 9,999 permutations). Identification of the nearest balance associated with clinical status was carried out as previously described (Suzuki et al. 2025), with flagellin clusters substituted for bacterial taxa. To account for repeated measurements across time points, each cross-validation iteration for nearest balance construction included a single randomly selected time point per subject (100 iterations). PERMANOVA was similarly applied across such iterations, and p-values from all runs were averaged to derive the final significance estimate.

## Results

### Human gut microbiome derived flagellins predicted to be stimulatory or silent: FlaPro annotations and model

Model performance was evaluated across 100 train/test splits. The resulting confusion matrix (Fig. 2) demonstrates reliable classification of flagellins into “stimulatory” and “silent” categories. The performance metrics indicate robust predictive capacity, with a specificity of 0.875, sensitivity of 0.83, and a test accuracy of 0.857 for the final model (Table 1). Random Forest feature importance analysis revealed that alignment-based features were the primary drivers of prediction accuracy (Suppl. Fig. 2). The consistently high performance across splits suggests the model effectively captures the key characteristics of the target phenotypes. Further validation through label randomization tests showed that predictive performance declined markedly when labels were randomized, indicating that the model learned genuine biological patterns rather than artifacts or biases (Suppl. Fig. 3 A,B).

**Table 1.** Model performance metrics summary

| **Metric** | **Value** | **Metric** | **Value** |
| --- | --- | --- | --- |
| **TestAccuracy** | 0.86 | **Balanced Accuracy** | 0.85 |
| **AccuracyPValue** | 0.001 | **Sensitivity** | 0.83 |
| **Kappa** | 0.71 | **Specificity** | 0.875 |
| **AccuracyLower** | 0.673 | **Pos Pred Value** | 0.83 |
| **AccuracyUpper** | 0.96 | **Neg Pred Value** | 0.875 |
| **AccuracyNull** | 0.57 | **Prevalence** | 0.43 |
| **TrainAccuracy** | 0.77 | **Detection Rate** | 0.36 |

**Figure 2.**
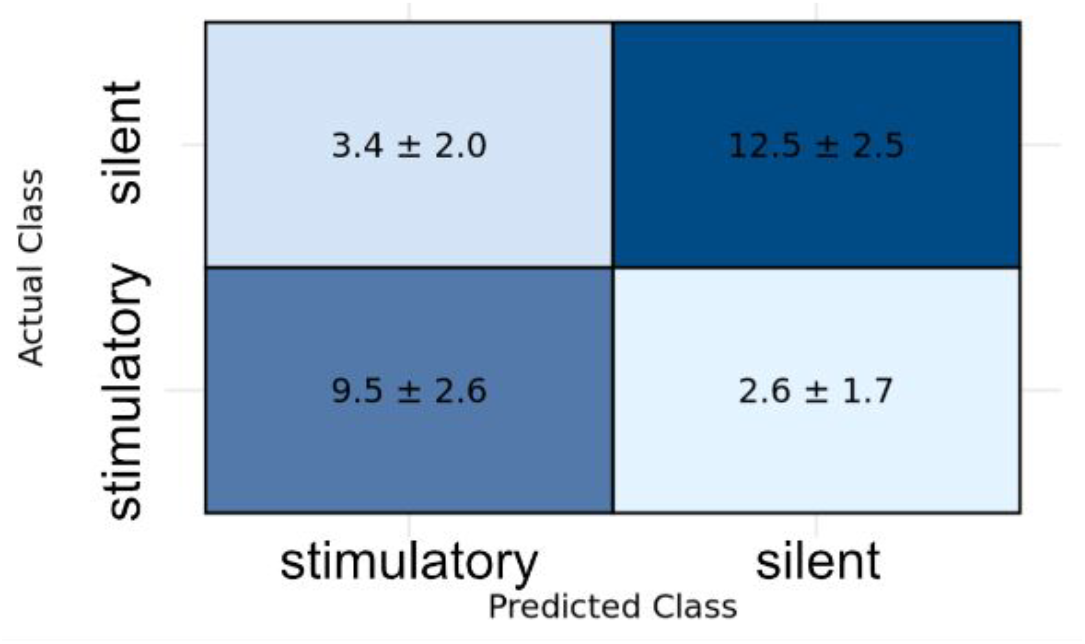
Performance evaluation of the classification model. Confusion matrix averaged over 100 randomized iterations (mean ± standard deviation).

Most of the cases that were misclassified in both training and test sets were located near the threshold separating the two phenotypes on the experimental TLR5 binding/activity plot (Suppl. Fig 4 A), and in these cases, the predicted class probabilities were relatively low. Notably, the farther a mismatch was from the experimental boundary, the lower the model’s confidence in its prediction, indicating that misclassifications often reflect underlying biological ambiguity rather than model error.

The experimental classification threshold is based on binding and activity assay values for the “FliC PIM” mutant variant, used as a negative control for binding at the TLR5 primary binding site (Yoon et al. 2012; Clasen et al. 2023). The experimental classification threshold is based on binding and activity assay values for the “FliC PIM” mutant variant, which is impaired at the primary interface (D1) involved in TLR5 recognition (Yoon et al. 2012; Clasen et al. 2023). However, as previously reported (Clasen et al. 2023), many flagellins display activity values close to this threshold. While strong binding at the primary interface can occur in both stimulatory and silent flagellins, signaling is believed to depend on an additional interaction at the D0 domain. This supports the idea that TLR5 interaction reflects a continuum rather than a binary trait – and suggests that the model captures this gradient to a meaningful extent.

Before applying the model to the full set of gut-derived flagellins, a small subset lacking key features was excluded, leaving 2,462 sequences for prediction. Among these, the majority were classified as stimulatory, with silent flagellins comprising approximately 15% of the set (Suppl. Fig. 4 B). We also observed that some CD-HIT clusters used for ShortBRED marker construction contained flagellins with differing predicted phenotypes (Suppl. Fig. 4 C). To address this, we assigned a consensus phenotype to each cluster when one phenotype predominated and the classification confidence score for alternative types remained low (close to 0). The cases with low classification confidence across all members should be interpreted with caution.

### Validating FlaPro on simulated data

To evaluate the accuracy of FlaPro in detecting and quantifying diverse flagellin gene sequences in metagenomes, we tested the FlaPro pipeline on a set of 6 simulated metagenomes (n=6) including an increasing number of known gut flagellin sequences (1 to 10). Preliminary results showed that ShortBRED Quantify reliably recovered the expected flagellin sequences and overall preserved their proportions (Suppl. Fig. 5)). While a large number of additional clusters (291 vs. 25 simulated) appeared in the profiles, their abundance values were 3-6 orders of magnitude lower than those of the expected features. Further evaluation using simulated datasets with real-life community complexity will provide additional insight into profiling accuracy.

### Linking gut flagellome to disease: IBD

Processing of a published microbiome dataset from IBD patients and healthy subjects (Lloyd-Price et al. 2019) in FlaPro showed that, among the 1,322 flagellin clusters in the reference database, only 22 (MGX) and 14 (MTX) were prevalent under our defined threshold (Suppl. Fig. 1 B,C; see Methods). In line with our earlier analysis (Clasen et al. 2023), experimentally tested flagellins comprised a small fraction of the detected flagellome (Suppl. Fig. 6 A). Silent flagellins were consistently less abundant than stimulatory ones – by approximately an order of magnitude – according to both experimental and predicted annotations (Suppl. Fig. 6 AB), with the stimulatory group being the primary contributors to the inter-sample variation (e.g., MTX: Fig. 3 A).

**Figure 3.**
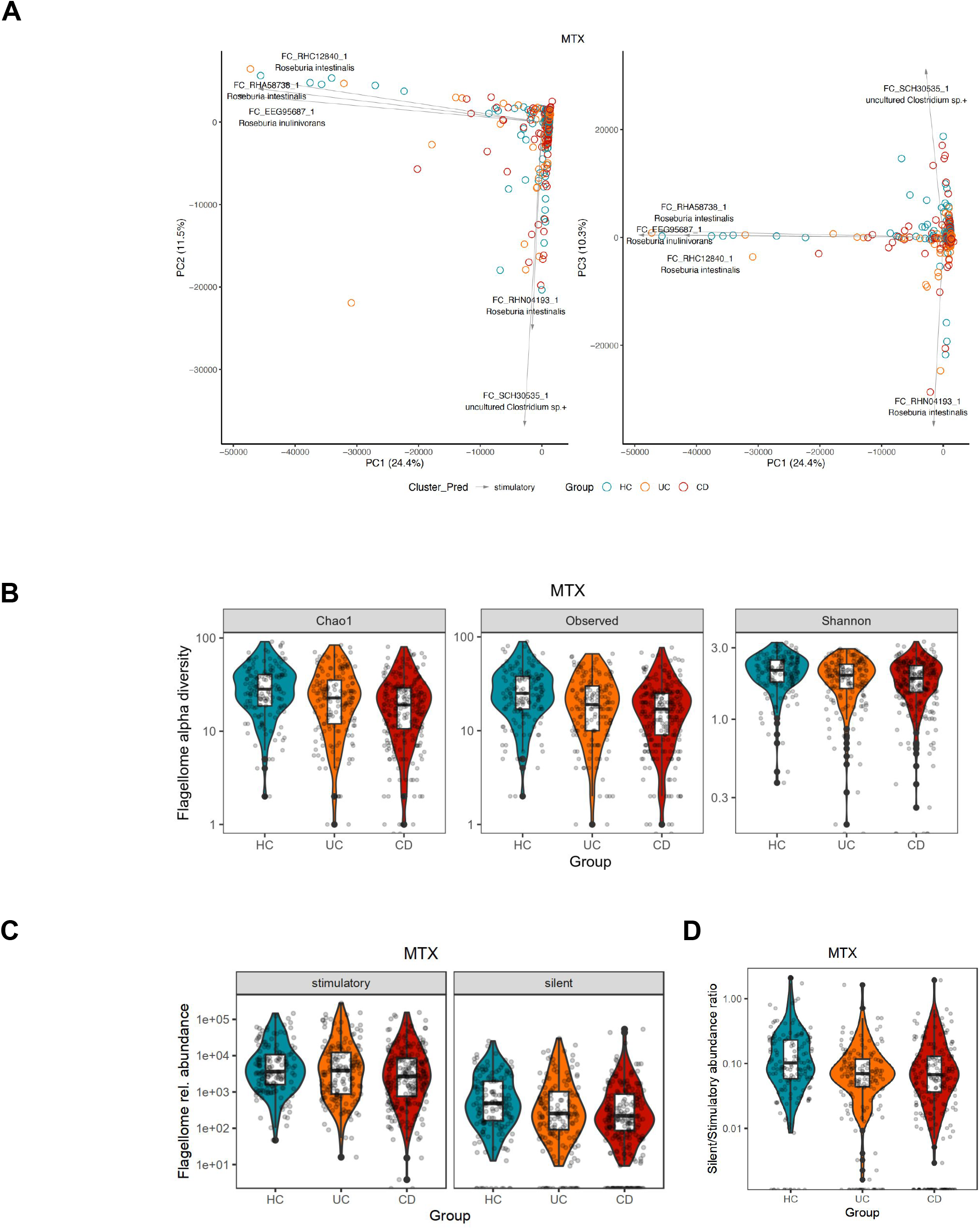
Altered gut flagellome expression in IBD. A) Principal coordinate analysis (PCoA) of MTX-derived flagellome profiles. Biplots show the first three components, with arrows indicating the direction of the top five contributing features. Based on Euclidean distances between flagellin relative abundance profiles. Fourteen statistical outliers (>3 s.d. from the mean) were excluded for clarity (final n = 562 samples). B) Reduced flagellome alpha diversity across IBD groups, measured by three indices. C) Lower total flagellin relative abundance in IBD. D) Decreased silent-to-stimulatory flagellin expression ratio. Each point represents one sample (multiple time points per subject are presented); distributions are shown as violin plots with log-scaled y-axes.

When applying alpha diversity metrics to the data, we observed IBD was associated with a reduced flagellome repertoire – at the level of encoded potential (MGX) as well as of the expression (MTX) (Fig. 3 B). This reduction was more pronounced in CD than in UC, particularly when comparing all three groups using the Chao1 metric in MTX (p = 4 × 10−^4^, linear mixed-effects model; see Methods). Stratified analysis showed this diversity loss affected both silent and stimulatory flagellins (Suppl. Fig. 7). However, it was only for the silent flagellins that we observed their total abundance reduced in both UC and CD compared to HC (for MGX and MTX/MGX) – in line with the decreasing silent/stimulatory abundance ratio (MGX and MTX; Fig. 3 C). Full statistical results are available in Suppl. Tables 2, 5.

Pathology-associated flagellome depletion was also evident at the individual cluster level. Differential analysis using the DiseaseScore variable in a three-group comparison showed that out of 22 major flagellin clusters, six (four stimulatory and two silent) were significantly decreased in abundance in MGX (adj. p < 0.05; Fig. 4A); at the level of expression, the respective number was 4 and for the MTX/MGX ratio – 2 (Suppl. Tables 3, 6, 8). More specifically, in CD patients relative to healthy controls, we observed reduced metagenomic abundance of flagellin clusters, primarily from commensal genera such as *Butyrivibrio* and *Agathobacter* (Suppl. Figure 8 A, Suppl. Table 9). MTX analysis showed decreased expression for 8 clusters, while the MTX/MGX ratio revealed reductions in 3 (Suppl. Figure 8 B,C; Suppl. Tables 10 and 11), suggesting the pathology-associated depletion of expressed flagellome occurs not just due to the decreased representation of the respective flagellated bacteria in the community but also due to the transcriptional downregulation within the respective taxa. In UC, although some associations yielded low raw p-values, none remained significant after correction for multiple testing – likely reflecting the smaller group size and, potentially, the milder clinical phenotype compared to CD.

**Figure 4.**
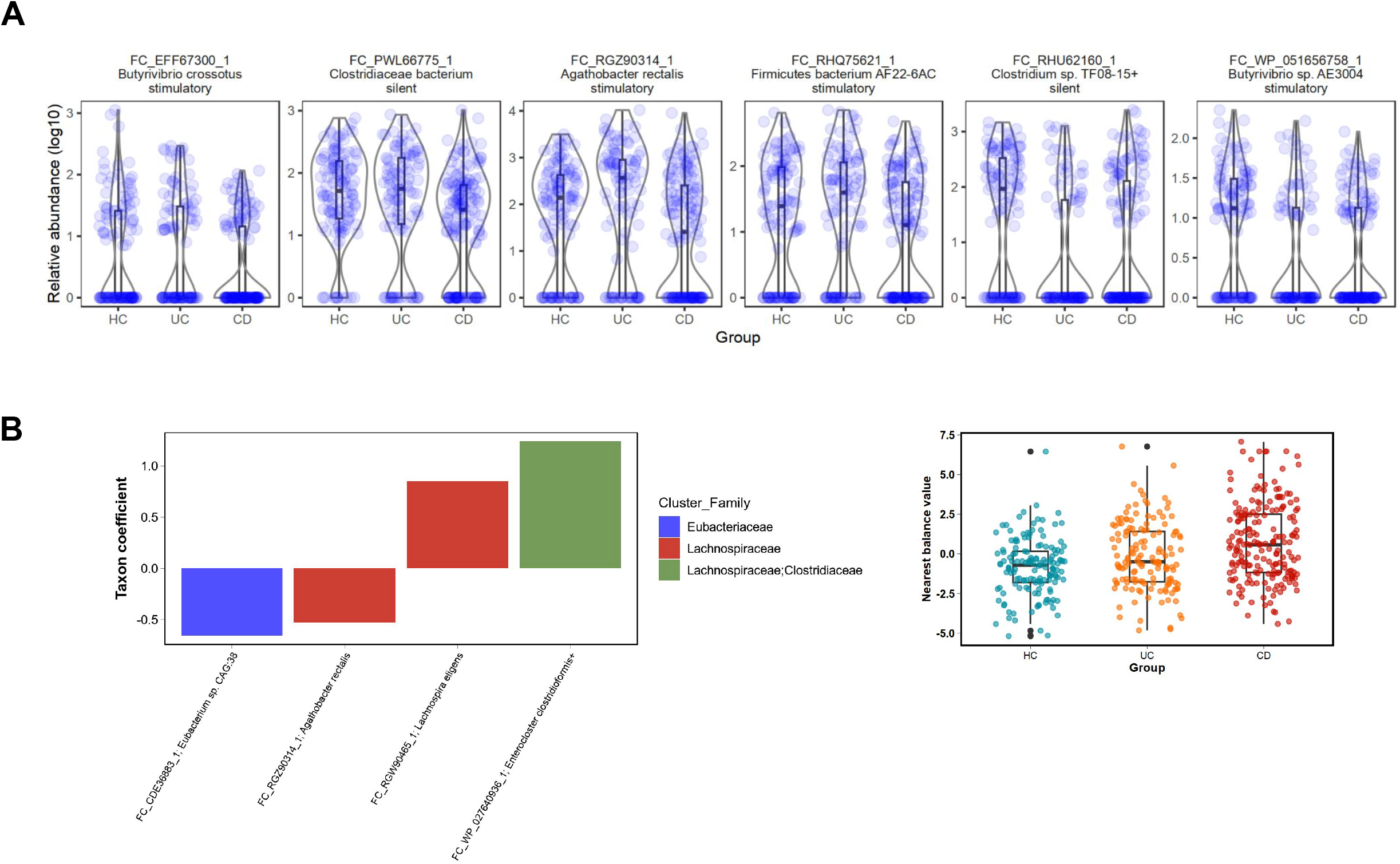
Differential analysis of flagellome-encoding potential in metagenomes. A) Component-based analysis across the clinical groups using a “disease score” variable (linear mixed-effects model on log-transformed data; adjusted p < 0.05). B) Compositional-aware analysis (CD vs. HC). Left: flagellin clusters with positive coefficients (numerator of the nearest balance) are enriched in CD as compared to HC, while those with the negative ones (denominator) are deplete. Right: balance values calculated for each metagenome using the centered log-ratio (clr)-transformed values, adjusted for age and sex; values for the UC subjects were calculated using the CD-vs-HC balance and are provided for reference.

From the alternative – compositional-aware – perspective, there was an overall tendency of link between the flagellome and IBD via MGX (DiseaseScore across all 3 groups: p = 0.06 ± 0.14, median ± s.d., repetitive-measures-aware PERMANOVA, see Methods), but not via MTX (median p > 0.2); this effect for MGX appeared driven primarily by CD rather than UC (Suppl. Table 12). For the CD vs. HC comparison in MGX, the nearest balance consisted of 2 flagellin clusters positively associated with the disease (numerator) and 2 negatively associated (denominator) (Fig. 4 B). We did not observe a clear enrichment of either silent or stimulatory classes in either part of the balance. Comparison of the component-based and compositional scenarios’ results suggests that the altered ratios between the distinct flagellins are less likely to play a role in IBD than their abundance relative to all genes of the bacterial community.

## Discussion and conclusions

Flagellin diversity is vast and this diversity has functional consequences with respect to host immune receptors. For the human innate immune receptor TLR5, flagellins can range how they tune the receptor response from highly stimulatory to silent. Here, we developed FlaPro to predict the TLR5 response of a flagellin from its sequence and other features. Cross-validation results suggest considerable performance of the prediction. We then integrated FlaPro into a workflow allowing the user to input MGX and MTX data for flagellin quantification, classification, and to understand the underlying dynamics of differences at the level of microbiome diversity versus expression patterns.

We demonstrated the utility of our FlaPro pipeline for profiling the flagellome and gaining insights into host–microbiome interactions using a large, publicly available multi-omics IBD gut microbiome dataset (Lloyd-Price et al. 2019). The alterations of the flagellome observed in this IBD dataset provide insight into how specific bacterial components may contribute to disrupted immune response to commensal microbiome. We hypothesize that the decrease of flagellome diversity and silent/stimulatory ratio may influence the repertoire and levels of anti-flagellin antibodies (Lodes et al. 2004; Alexander et al. 2021; Shen et al. 2008). Investigation of larger sample sizes with consideration of IBD clinical subtypes and severity, particularly in treatment-naive patients, will provide a more detailed understanding of how the flagellin-encoding potential and expression vary in these diseases.

The downstream secondary data analysis with an R notebook generator functionality can be flexibly adjusted to most common study design schemes. The raw features produced by the primary analysis can be readily input to feature engineering routines (for example, co-occurrence-based clustering) and more elaborate statistical approaches to differential abundance analysis (including those accounting for sparsity in data).

Our annotation strategy combining structure- and sequence-based prediction of function is extendable beyond human TLR5 to investigate flagellin interactions in other model animals (such as mice with their divergent TLR5 structure), in diverse niches (in environmental microbiome or plant microbiome). More broadly, this approach can be generalized to study other bacterial protein families that interact with host receptors, facilitating comprehensive, receptor-specific analyses of host–microbiome dynamics across a range of biological contexts.

Our methodology is not free from limitations. Firstly, a few of them are related to the set of the experimentally evaluated flagellins: it is relatively small, had been constructed with a bias towards healthy subjects’ microbiome and has underrepresented evasive phenotype due to the lack of known cases to date. We are currently conducting additional experiments to expand this training set and improve the model performance and generalizability. Secondly, in the current implementation of the feature construction, some flagellin clusters turn out to be mixed (by combining silent and stimulatory flagellins) thus complicating further interpretation of these features. In the IBD dataset analyzed here, the mixed clusters were low abundant and not associated with the clinical status. However, if profiling of future datasets produces associations for such features, their TLR5-related phenotypes can be elucidated experimentally. Additionally, as flagellins constitute a single protein with often very similar amino acid sequences, there is a risk of false positive assignments, especially when analyzing short-read sequences. Once a sufficient volume of long-read microbiome sequencing datasets becomes available, our pipeline can be updated to enable processing of the long reads to increase the profiling accuracy.

We anticipate that profiling of metagenomes from patients with diverse immune-related disorders using FlaPro will shed more light onto the contribution of gut microbiome to their pathogenesis. Future profiling of large metagenomic and metatranscriptomic datasets using FlaPro will help define the landscape of flagellome in health and uncover both condition-specific and shared alterations across diseases.

## Author contributions

AVT and REL conceptualized the study. AVT provided supervision and project administration. ABG optimized the alignment parameters and developed prototype scripts for flagellome profiling. AAB developed the flagellome profiling Snakemake pipeline, prepared the reference datasets, developed and evaluated the prediction model, performed predictions for the reference database, profiled the metagenomes. AVT developed the secondary analysis notebooks and processed the flagellomes. AAB and AVT wrote the manuscript. AVT, AAB, REL, and ABG revised the manuscript.

## Acknowledgements

We thank Miriam Haag, Michael Bell, James Marsh, and Elizaveta Kulaeva for their valuable input and insightful discussions throughout the development of this work.

## Notes

https://github.com/leylabmpi/FlaPro

### Competing Interest Statement

The authors have declared no competing interest.

