## Supplemental figures for "Human gut flagellome profiling using FlaPro reveals TLR5-related phenotype-specific alterations in IBD"

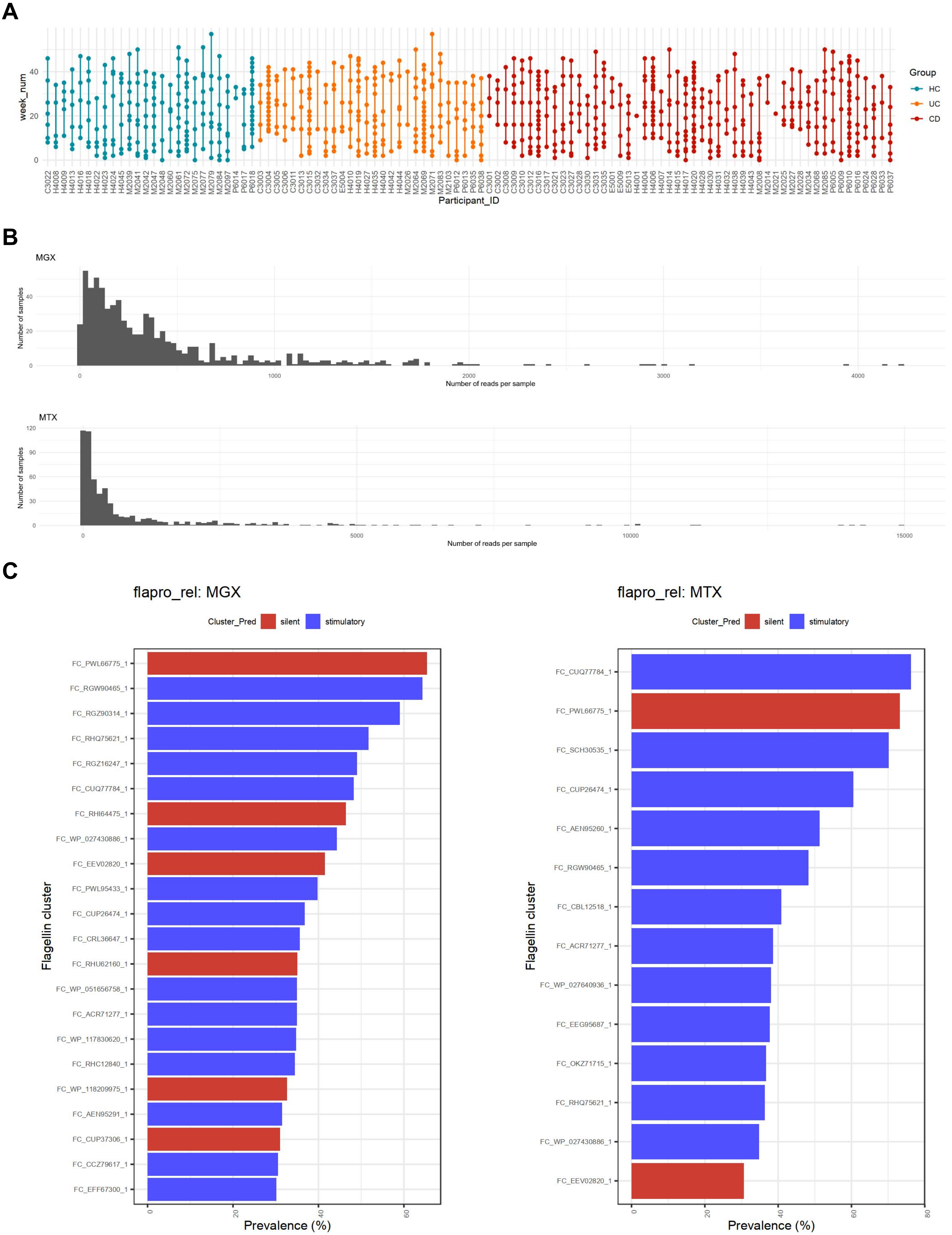


**Supplementary Figure 1. Overview of the IBD flagellome dataset.** A) Distribution of available time points per subject. B) Number of reads mapped to the flagellin reference set, separated by MGX and MTX layers. C) Prevalent flagellin clusters (prevalence >30%) with their predicted phenotypes.

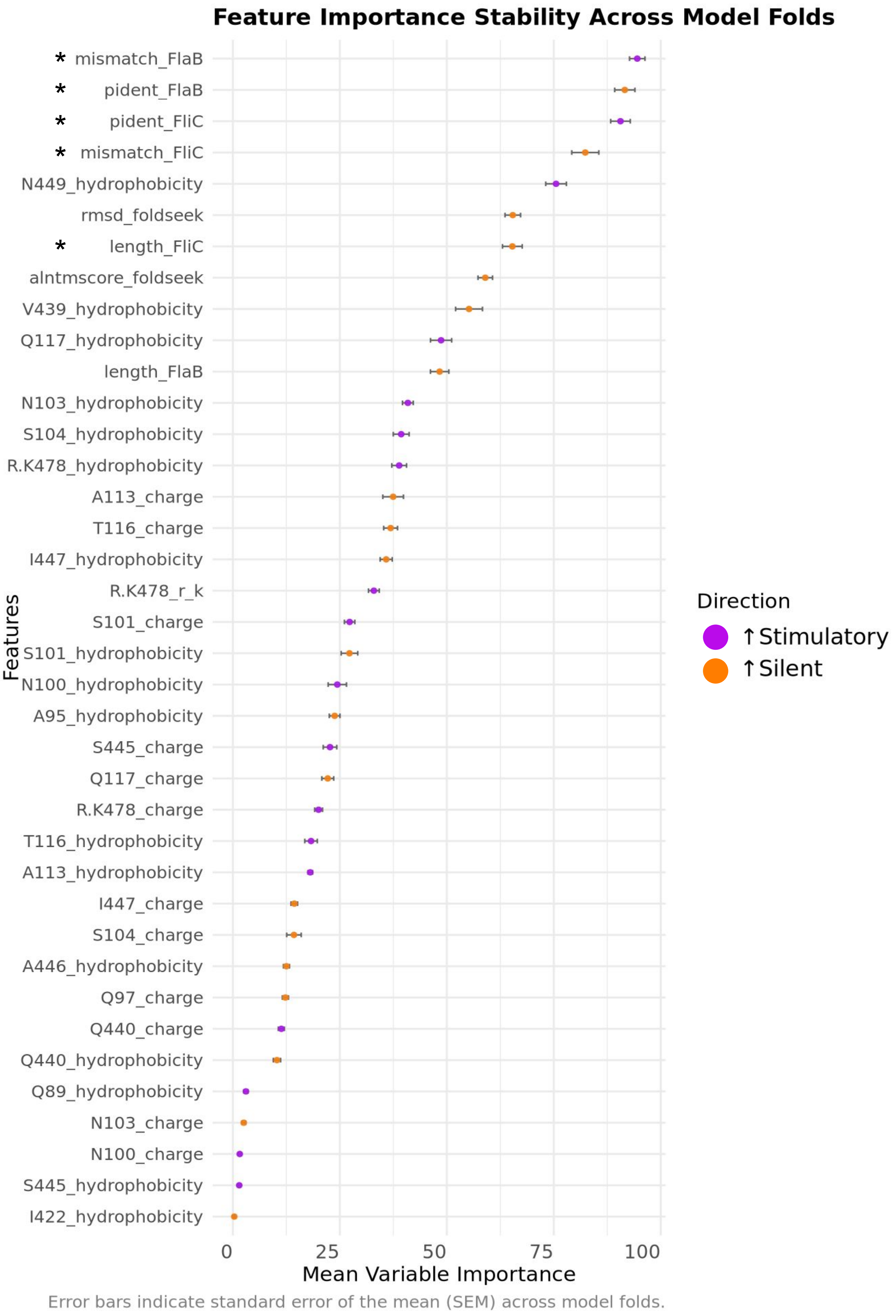


**Supplementary Figure 2. Feature importance in the classification model.** Mean variable importance across model folds; error bars indicate standard error of the mean (SEM). Color indicates direction of association (stimulatory vs. silent). Asterisks (*) mark high-impact sequence alignment-based features.

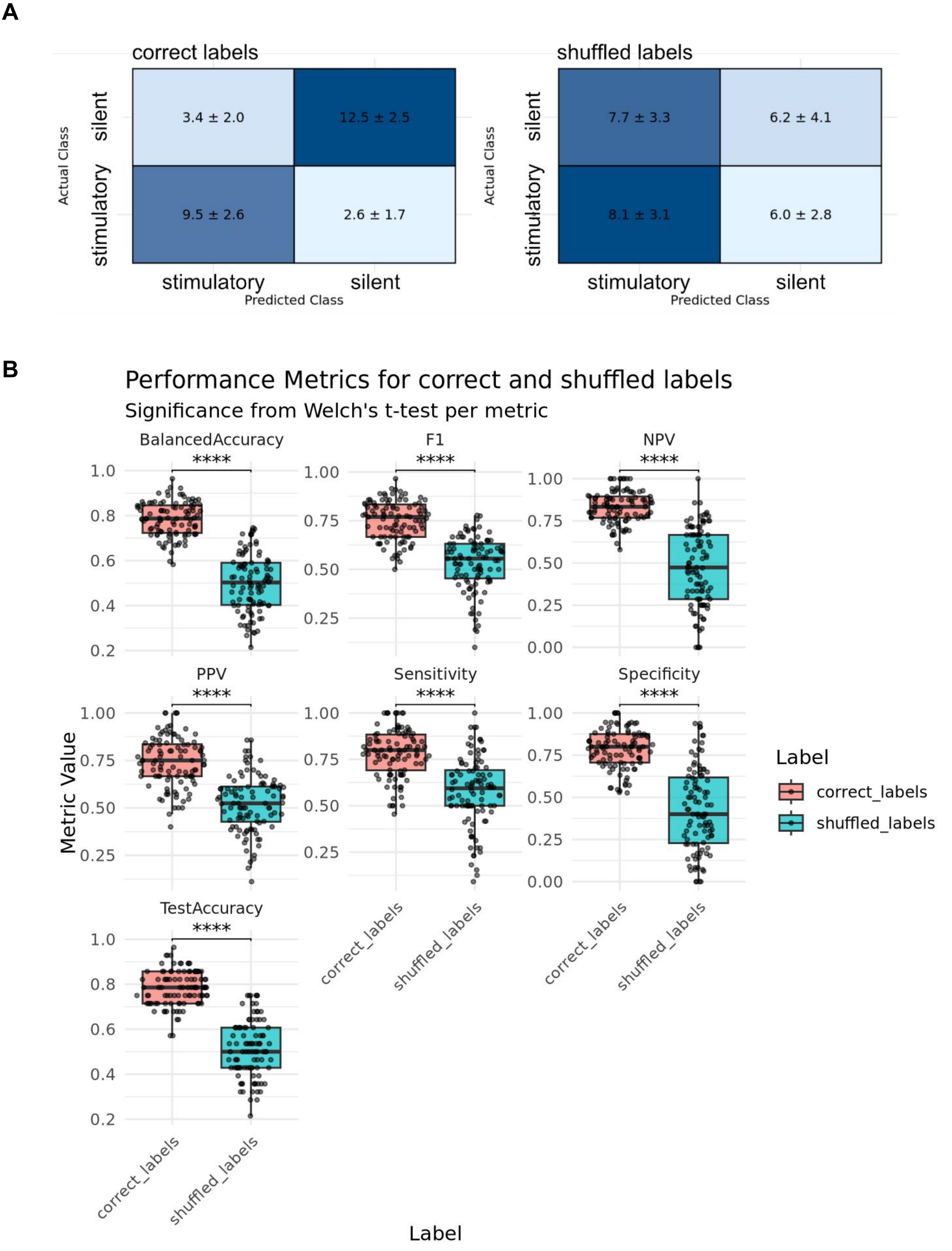


**Supplementary Figure 3. Label shuffling test across 100 iterations.** A) Confusion matrices for correct (left) and shuffled (right) labels; mean values ± SEM. B) Distribution of model quality metrics for true (coral) and shuffled (blue) labels. Significance assessed via two-sided t-test (**** $p < 0.0001$).

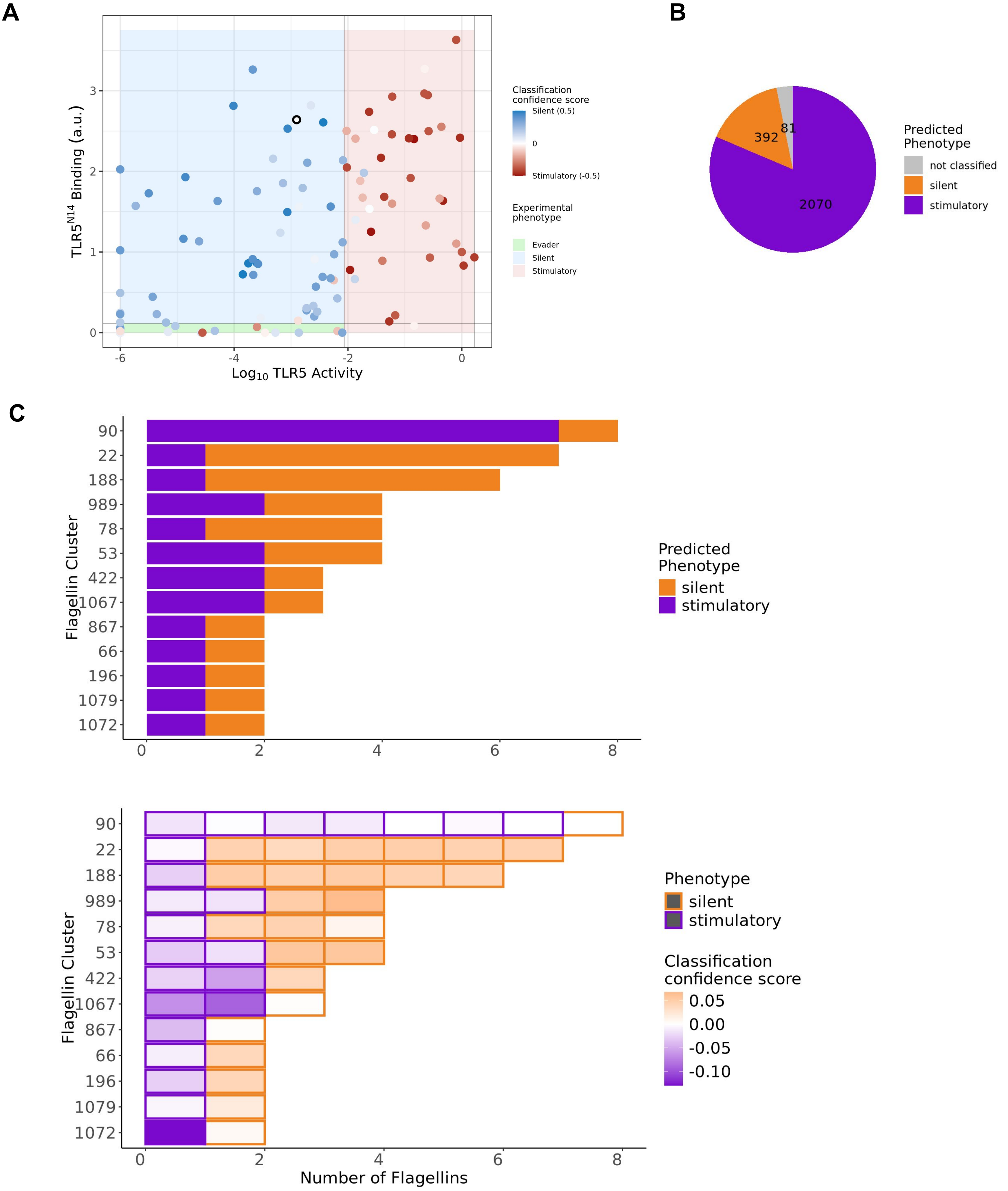


**Supplementary Figure 4. Predicted flagellin phenotypes.** A) TLR5 binding/activation plot based on Clasen et al, 2023 data with experimentally defined phenotypes shown as color-filled background regions and predicted phenotypes' classification confidence score overlaid as gradient-colored points (red: stimulatory, blue: silent). Dots with a black border reflect low-confidence classification (Sconfidence = 0). B) Proportions of predicted silent, stimulatory, and unclassified flagellins in the human gut microbiome derived database. C) Distribution of mixed phenotype clusters (as used by ShortBRED *quantify* module). Top: number of clusters with conflicting phenotype assignments. Bottom: prediction probabilities indicating confidence of phenotype assignment for mixed clusters.

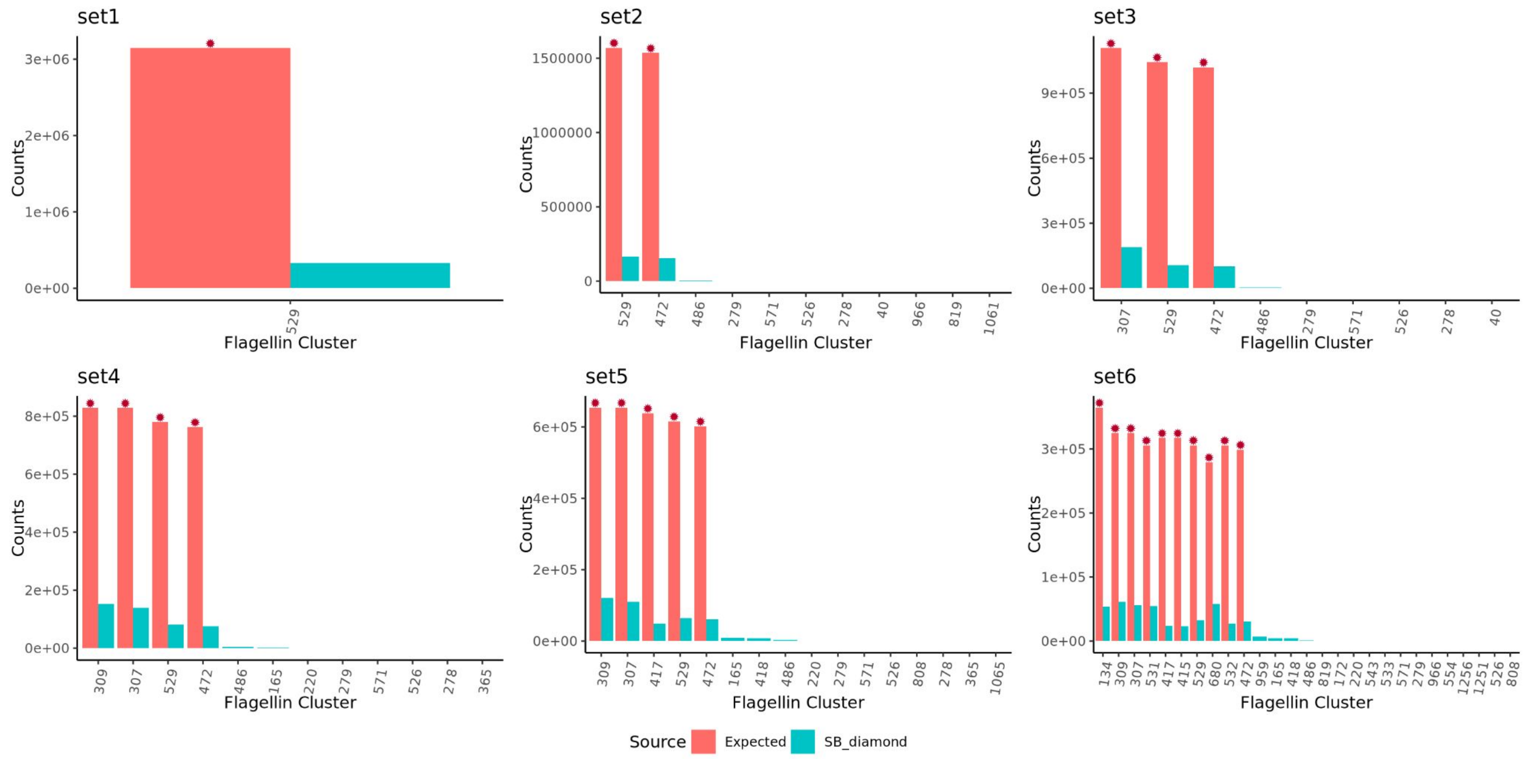


**Supplementary Figure 5. Validation on simulated data.** Coral bars (with stars) indicate ground-truth flagellin clusters; blue bars represent those detected by the FlaPro via ShortBRED *quantify* with DIAMOND alignment. Low-abundance features (<30 reads) are not shown.

A

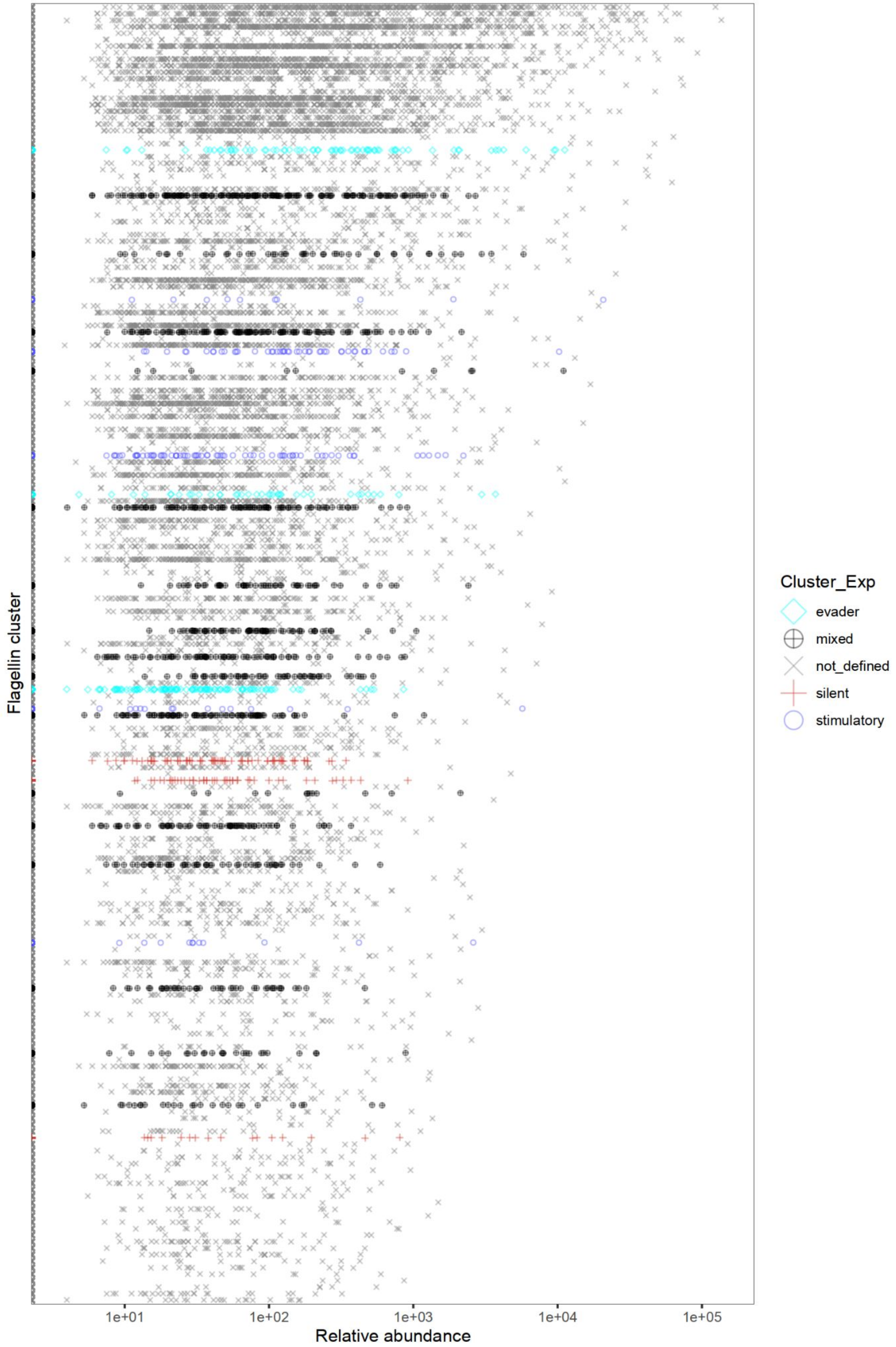


**Supplementary Figure 6. Most abundant flagellin clusters in the IBDMDB dataset.** Ranking of the top 200 clusters by mean relative abundance in MTX data. A) Annotated based on experimental data. B) Annotated using predicted phenotypes.

**B**

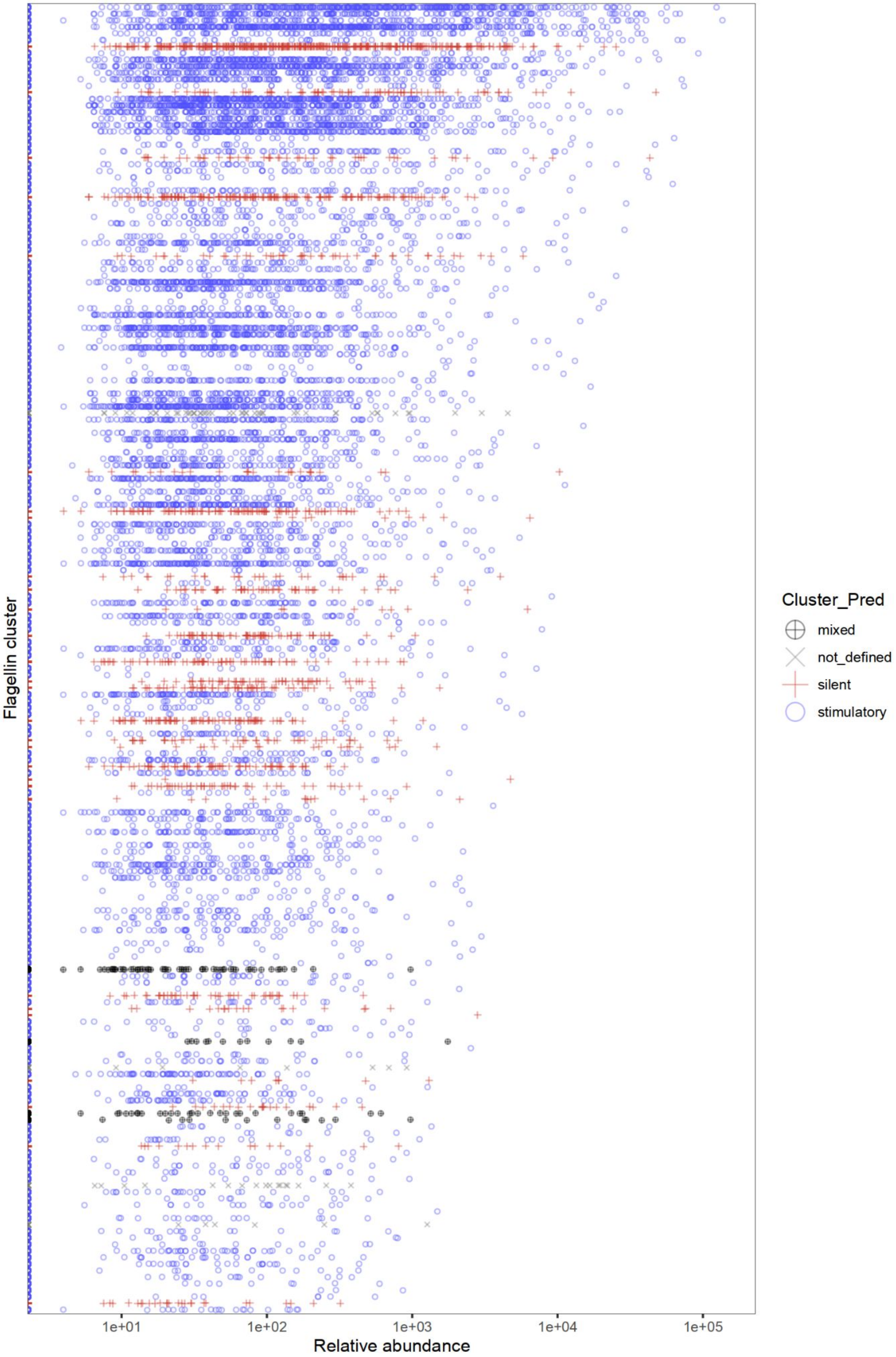
Flagellin cluster
Relative abundance
1e+01
1e+02
1e+03
1e+04
1e+05
Cluster_Pred
mixed
not_defined
silent
stimulatory

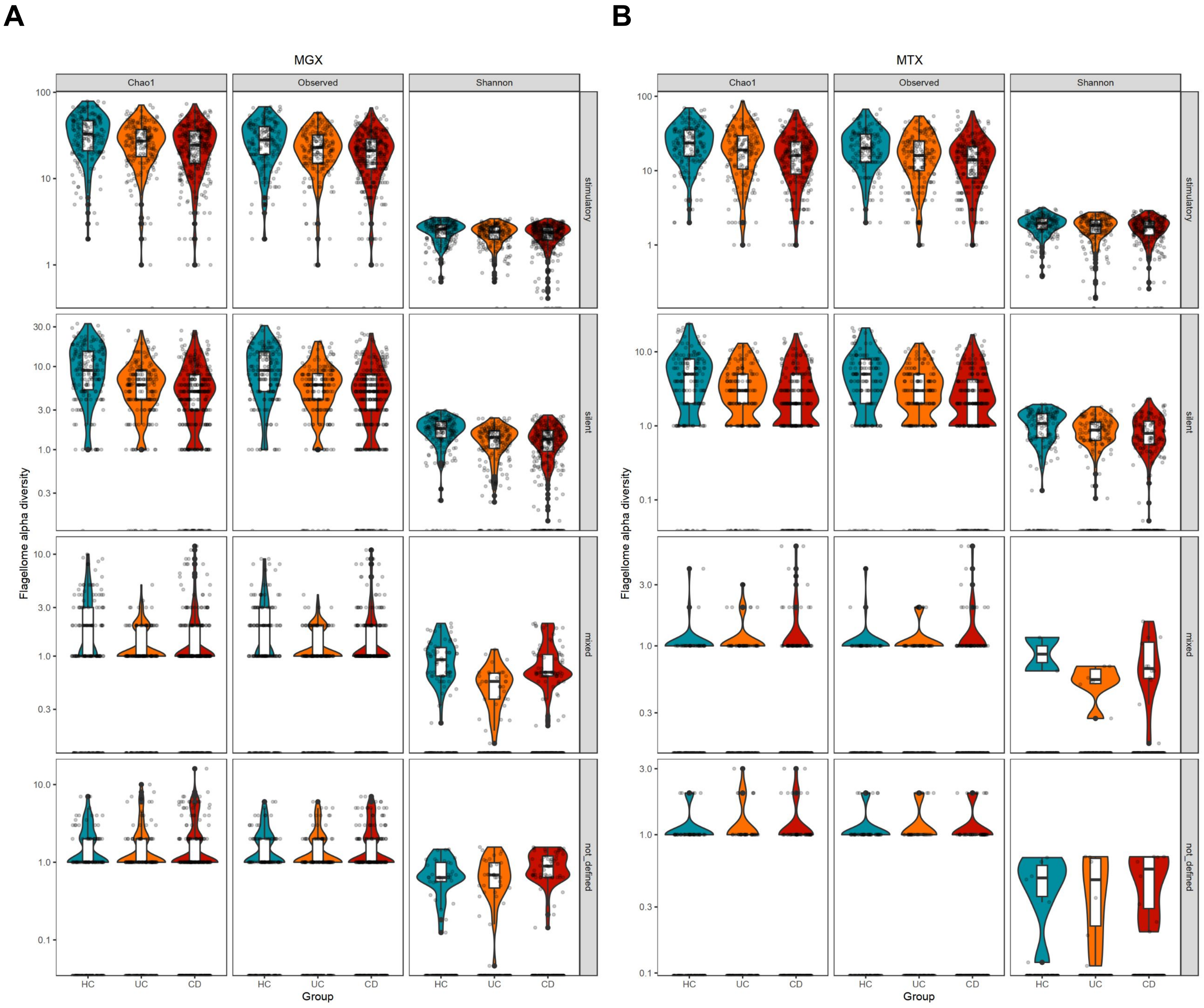


**Supplementary Figure 7. Flagellome alpha diversity stratified by flagellin class.** A) MGX data. B) MTX data. Diversity metrics stratified by flagellin phenotype (stimulatory, silent, mixed, undefined).

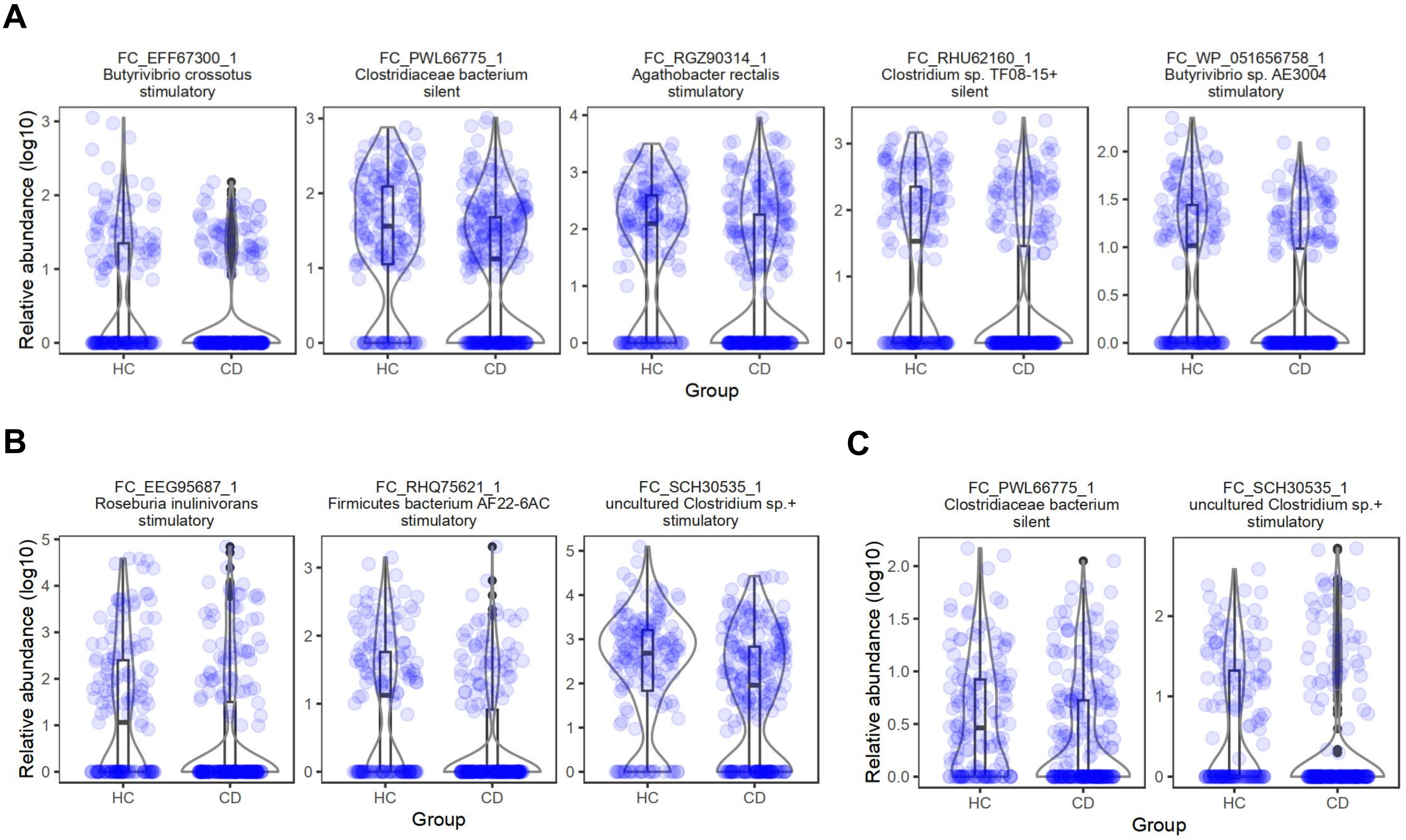


**Supplementary Figure 8. Differential abundance of flagellin clusters in CD vs. HC.** Component-based analysis across the prevalent flagellin clusters for: A) MGX, B) MTX, and C) MTX/MGX ratio. Linear mixed-effects model applied to log-transformed values (adjusted $p < 0.05$). "+" indicates clusters with multiple taxa (only the first taxon shown; full list in corresponding supplementary table). Point color indicates direction of change (blue = decreased in CD; red = increased).
